# A simple immunohistochemical method for perinatal mammalian ovaries revealed different kinetics of oocyte apoptosis caused by DNA damage and asynapsis

**DOI:** 10.1101/2024.09.05.611563

**Authors:** Hiroshi Kogo, Akiko Iizuka-Kogo, Hanako Yamamoto, Maiko Ikezawa, Yukiko Tajika, Toshiyuki Matsuzaki

## Abstract

Oocytes having meiotic defects are assumed to be eliminated by apoptosis in perinatal period. However, the oocyte apoptosis caused by meiotic defects has not been well analyzed, partly due to the great technical demands for tissue sectioning of perinatal ovaries. In the present study, we applied a squash method for immunohistochemical analysis of perinatal mouse ovaries as a substitute for tissue sectioning. As a result, we could show different kinetics of apoptosis caused by DMC1- and SPO11-deficiencies, indicating that DNA damage-induced apoptosis precedes asynapsis-induced apoptosis in mouse oocytes. Double mutant analysis revealed that only asynapsis-induced apoptosis was significantly dependent on HORMAD2. The present method is simple, easy, and able to analyze a sufficient number of oocytes for detecting infrequent events in a single specimen, accelerating detailed immunohistochemical analyses of mammalian ovaries during the fetal and perinatal periods.

## Introduction

In many mammalian species, the maximum number of female germ cells is set before birth, in contrast to the continuous production of male germ cells throughout life. In mice, the number of oocytes peaks at the onset of meiosis in the fetal ovary (∼15,000), and about one-half or two-thirds of oocytes die before the primordial follicle reserve is established just after birth (Morita and Tilly 1999; Findlay et al. 2015; Kaur and Kurokawa 2023). It has been well demonstrated that extensive oocyte loss occurs at 1-3 days post-partum (dpp) concomitant with germ cell cyst breakdown (Pepling and Spradling 2001; Bristol-Gould et al. 2006). This oocyte loss is likely a physiological process of selecting the surviving oocytes from the so-called nursing oocytes in the cysts (Pepling 2006; Niu and Spradling 2022) and thus is not a mechanism for eliminating defective oocytes.

Oocytes having meiotic defects are also known to be eliminated in the perinatal period, as revealed by gene-targeting studies of meiosis-essential genes in mice (Bolcun-Filas and Schimenti 2012; Subramanian and Hochwagen 2014). For example, SPO11 is the enzyme that catalyzes DNA double-strand breaks (DSBs) required for meiotic recombination, and its deficiency results in extensive synapsis failure (Baudat et al. 2000). DMC1 is the meiosis-specific recombinase, and its deficiency causes many unrepaired DSBs in addition to extensive synapsis failure (Pittman et al. 1998). The oocyte loss occurred in both SPO11- and DMC1-deficient mouse ovaries and was more severe in the latter than the former, suggesting that meiotic prophase checkpoint(s) respond to both DNA damage and synapsis failure (Di Giacomo et al. 2005). Our and other previous studies showed that HORMAD2, which is recruited to unsynapsed axes of the synaptonemal complex by HORMAD1, plays an essential role in the meiotic prophase checkpoint(s) (Kogo et al. 2012; Wojtasz et al. 2012). HORMAD2 deficiency rescues SPO11-deficient oocytes almost completely (Kogo et al. 2012; Wojtasz et al. 2012) but only partially DMC1-deficient oocytes (Rinaldi et al. 2017), suggesting a different contribution of HORMAD2 in DNA damage and asynapsis checking (Ravindranathan et al. 2022). These observations are mainly based on the analysis of survived oocytes, and the apoptosis itself driven by the meiotic prophase checkpoint(s) has yet to be well demonstrated.

To analyze oocyte apoptosis and other events immunohistochemically, tissue sectioning techniques have been one of the best approaches (Pepling and Spradling 2001; Rodrigues et al. 2009; Niu and Spradling 2022). However, the preparation procedures are time-consuming, and sectioning small tissues like perinatal ovaries is technically difficult. Furthermore, it would be necessary to immunostain the more sections to detect and analyze certain events the more comprehensively, when tissue sections were used for analyses (McClellan et al. 2003).

In the present study, we applied a modified squash method to analyze oocyte apoptosis in perinatal mouse ovaries. We could successfully analyze a sufficient number of oocytes to reveal the different kinetics between DNA damage- and asynapsis-caused oocyte apoptosis, proving the present method suitable for analyzing infrequent oocyte events immunohistochemically.

## Materials and Methods

### Mice

The *Hormad2-*targeted mice were generated as described (Kogo et al. 2012). The *Spo11*-targeted mice were kindly provided by S. Keeney and M. Jasin. The *Dmc1*-targeted mice were kindly provided by J. C. Schimenti. Mice were bred and maintained under specific pathogen-free conditions in accordance with the Animal Care and Experimentation Committee at Gunma University (approval no. 17-017 and 22-017). The genotypes of *Spo11* and *Hormad2* alleles were determined by PCR as described (Baudat et al. 2000; Kogo et al. 2012). The genotype of *Dmc1* allele was determined by PCR as described (Pittman et al. 1998) or using the forward primer 5’-GCCACTAAGGATCAGACAAG-3’ and the reverse primer 5’-GACGTTGACAGAGTTTTGAC-3’ that amplify 468 bp and 598 bp fragments from wild-type (B6-derived) and mutant (129-derived) alleles, respectively.

### Simplified squash method for perinatal mouse ovary

A previously described squash technique for mouse seminiferous tubules (Page et al. 1998) were applied for perinatal mouse ovaries with some simplifications and modifications. Perinatal mouse ovaries were excised from fetal 17- and 18-days post coitus (dpc) and neonatal 0- and 1-days post partum (dpp) female mice under a dissecting microscope. Ovaries were rinsed with PBS in a tube and then transferred into 200 μl of fixative solution (2% paraformaldehyde in PBS containing 0.05% Triton X-100) by a micropipette set at 10 μl. After a 5 -10 min fixation, a single ovary was transferred on a silane-coated slide with 10 μl fixative by a micropipette. A 24 mm x 24 mm coverslip was put on the ovary, which was squashed by thumb pushing and quickly frozen in liquid N_2_. These procedures are schematically visualized in Figure 1.

**Figure 1.**
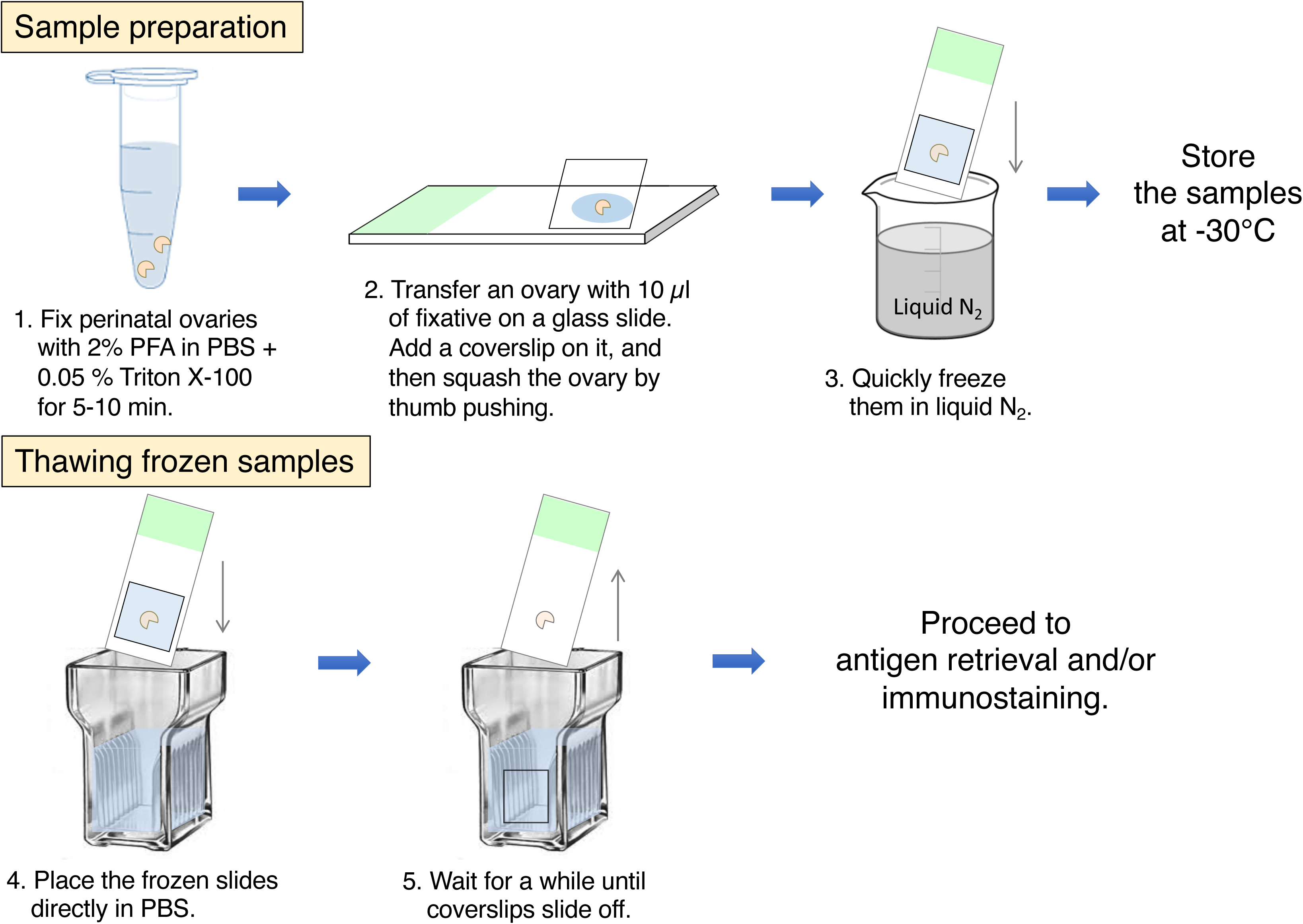
Schematic representation of squash method. Upper panel (Sample preparation) shows the fixation, squashing, and freezing steps before storage in a freezer. Lower panel (Thawing frozen samples) shows how to remove coverslips with sample thawing before antigen retrieval and/or immunostaining.

### Antigen retrieval and immunofluorescent staining

For antigen retrieval, squashed specimens were placed in DAKO Real Target Retrieval Solution (pH 6) (S2031; DAKO) and heated in a microwave oven (MI-77; Azumaya) for 20 min at 98°C. After cooling down and washing with PBS, specimens were incubated with 10% normal donkey serum/PBS to block non-specific binding of antibodies. Immunofluorescent staining was performed using the following primary antibodies diluted in PBS: guinea pig anti-SYCP3 antiserum (1:6,000) and rabbit anti-cleaved PARP1 (1:200, Cell Signaling Technology #9544). The following secondary antibodies were used for detection: Alexa 488-conjugated donkey anti-guinea pig IgG (1:200) and Alexa 594-conjugated donkey anti-rabbit IgG (1:1000).

### Microscopy and image analyses

Images were captured from the entire area of each ovary containing SYCP3-positive oocytes using the 40x objective lens of a BX62 microscope equipped with epifluorescence optics (Olympus), and a CoolSNAP K4 CCD camera (Photometrics) controlled by MetaMorph software version 6.1 (Molecular Devices). The number of SYCP3- and cleaved PARP1-positive nuclei was manually counted with genotypes blinded to the counters using the Photoshop counting tool (Adobe).

### Statistics

Statistical analyses were performed using EZR software (Kanda 2013). To compare frequencies of oocyte apoptosis in *Spo11*^-/-^ and *Dmc1*^-/-^ ovaries with that in wild-type ovary at each age (Fig. 2D), we performed the non-parametrical Kruskal-Wallis test followed by post hoc Steel’s test. To compare frequencies of oocyte apoptosis in wild-type, *Spo11*^-/-^, and *Dmc1*^-/-^ ovaries with those in corresponding *Hormad2*^-/-^ ovaries at each age (Fig. 3), we performed the Mann-Whitney U test. A *P* value of < 0.05 was considered statistically significant.

**Figure 2.**
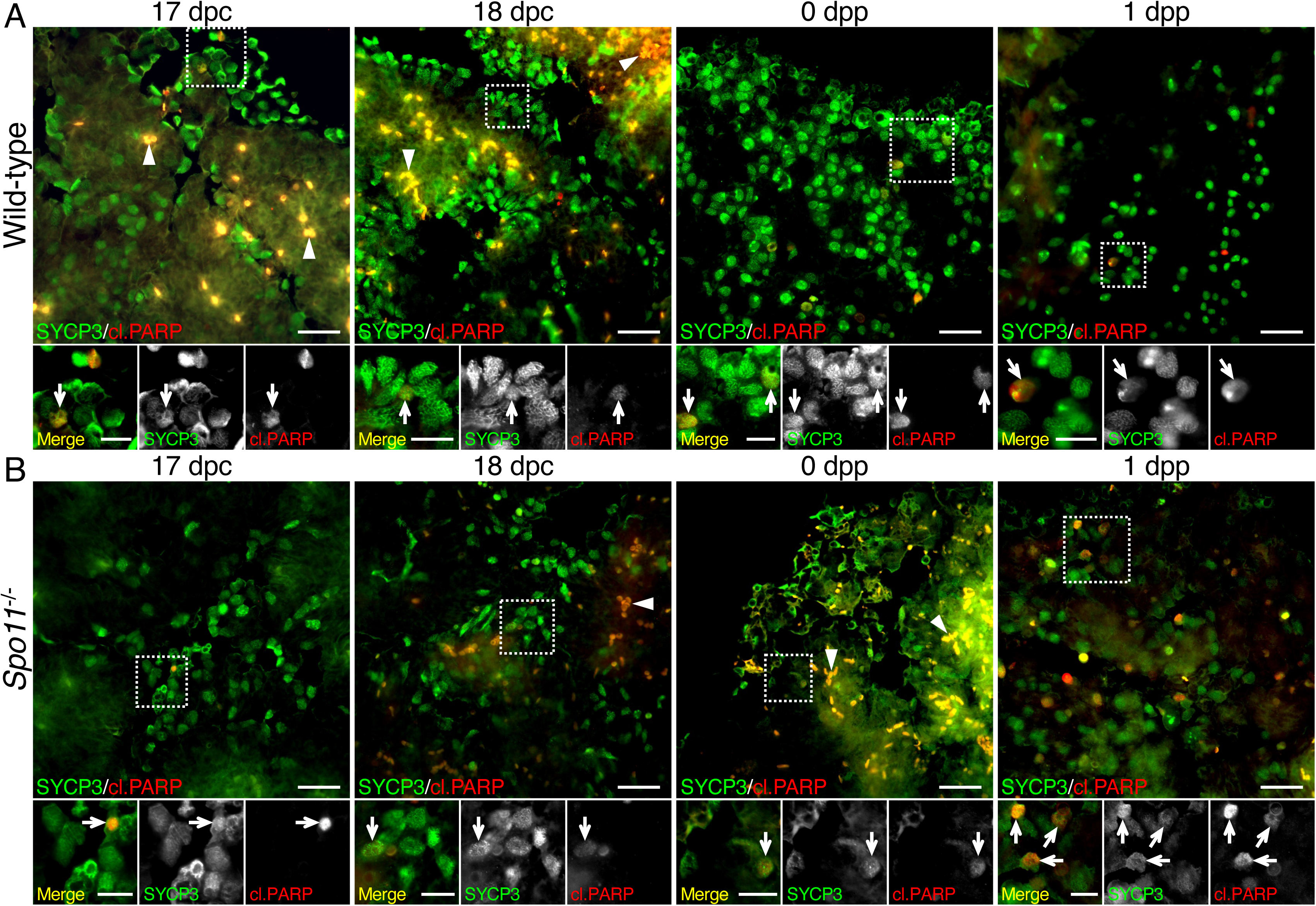
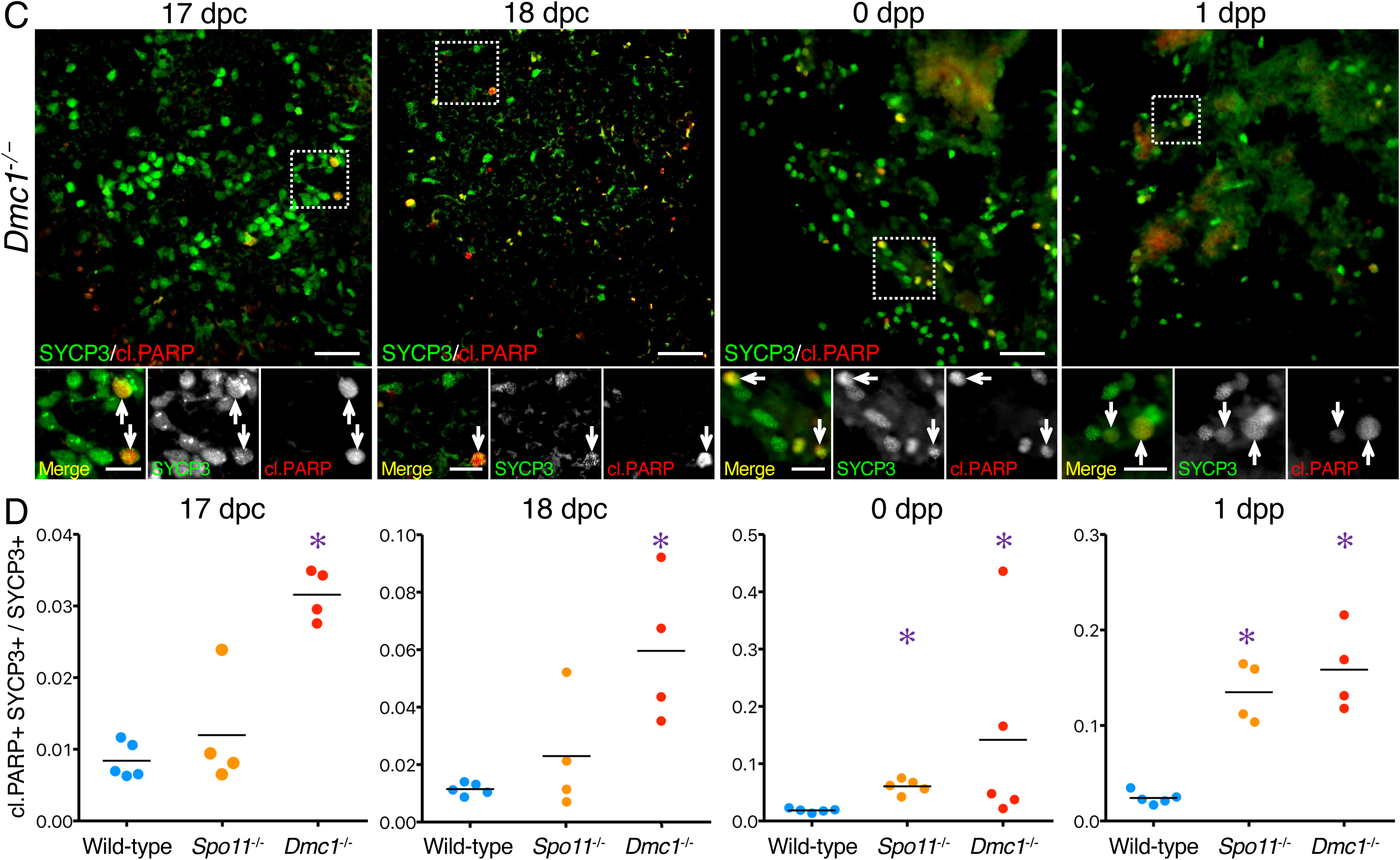
Immunohistochemical staining of oocyte apoptosis in squashed ovaries. Representative immunofluorescent images of wild-type (A), *Spo11*^-/-^ (B), and *Dmc1*^-/-^ (C) perinatal mouse ovaries. SYCP3 (green) and cleaved PARP1 (red) visualize meiotic oocyte nuclei and apoptotic nuclei, respectively. Squared areas are shown with a magnified view in merged and monochrome images of each staining below. White arrows indicate cleaved PARP1-positive apoptotic nuclei in magnified views. White arrowheads indicate erythrocytes showing autofluorescence. Results are representative of 4-5 independent experiments using different animals. Scale bars are 50 and 20 µm in full and magnified views, respectively. (D) Scatter dot plots for the frequency of apoptotic oocyte nuclei in wild-type, *Spo11*^-/-^, and *Dmc1*^-/-^ perinatal mouse ovaries. Bars indicate the mean of frequency. Statistical analysis was performed using the Kruskal-Wallis test, followed by the Steel’s test to compare the mutant groups *(Spo11*^-/-^ and *Dmc1*^-/-^) with the control group (wild-type). * p < 0.05.

**Figure 3.**
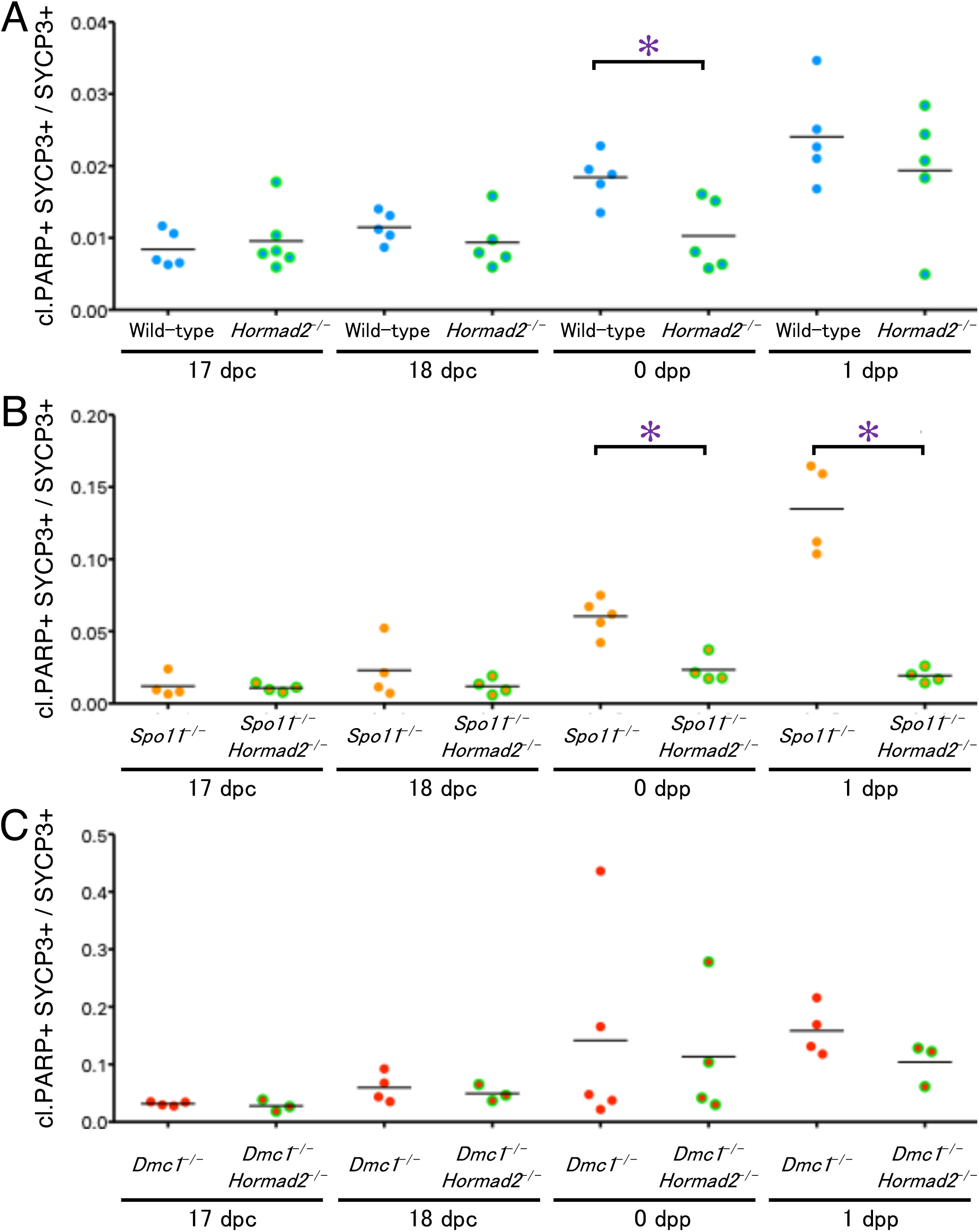
HORMAD2-dependency of oocyte apoptosis in wild-type, *Spo11*^-/-^, and *Dmc1*^-/-^ ovaries. Scatter dot plots for the frequency of apoptotic oocyte nuclei in wild-type and *Hormad2*^-/-^ (A), *Spo11*^-/-^ and *Spo11*^-/-^ *Hormad2*^-/-^ (B), and *Dmc1*^-/-^ and *Dmc1*^-/-^ *Hormad2*^-/-^ (C) perinatal mouse ovaries. The wild-type and single mutant ovaries’ data were the same as those in Fig. 2D. Bars indicate the mean of frequency. Statistical analysis was performed using the Mann-Whitney U test to compare oocyte apoptosis frequency between the presence and absence of HORMAD2 at each genotype and age. * p < 0.05.

## Results and Discussion

### Simple preparation of perinatal mouse ovaries for immunofluorescence

Squash preparation of mouse meiocytes was previously reported as a method for minced seminiferous tubules (Page et al. 1998). We applied this method for perinatal mouse ovaries with some simplifications and modifications. The procedures are summarized in Figure 1. This preparation is rapid and simple, having no technical difficulty compared to embedding and sectioning small tissues such as perinatal mouse ovaries. The coverslips practically did not peel off ovarian tissues without any coating, such as siliconization, which was confirmed by staining the coverslips with DAPI (Fig. S1A). Importantly, we found that heat-induced antigen retrieval techniques are applicable to the squash samples, enabling a wide variety of antibodies to be used for immunostaining. The present study used synaptonemal complex protein SYCP3 as a marker of meiotic chromosome axes, whose characteristic filamentous staining is easily distinguishable from diffuse non-specific staining (Fig. S2). As an apoptosis marker, cleaved PARP1, which shows a nuclear localization, was employed to identify apoptotic oocytes by colocalization with SYCP3-positive nuclei. Meiotic and apoptotic oocytes were counted manually, and genotypes were blinded to the counters. Some SYCP3-positive structures were omitted when diffusely stained (Fig. S2, white and yellow arrowheads) or severely deformed (Fig. S2, arrows). As a result, we could analyze about 700∼1,700 oocytes on average and up to about 4,000 oocytes from a single wild-type ovary (Table 1), where 5,000∼10,000 oocytes were estimated to exist (Pepling and Spradling 2001; Morohaku et al. 2017). This result shows that the present simple method can analyze a sufficient number of oocytes on a single specimen of perinatal mouse ovaries.

**Table 1.** Maximum, minimum, and average numbers of meiotic oocytes analyzed for each genotype and age group.

|  | 17 dpc |  |  | 18 dpc |  |  | 0 dpp |  |  | 1 dpp |  |  |
| --- | --- | --- | --- | --- | --- | --- | --- | --- | --- | --- | --- | --- |
|  | max. | min. | avr. (n) | max. | min. | avr. (n) | max. | min. | avr. (n) | max. | min. | avr. (n) |
| Wild-type | 2160 | 687 | 1451.0 (n=5) | 4152 | 268 | 1677.4 (n=5) | 1807 | 286 | 813.6 (n=5) | 1607 | 221 | 691.4 (n=5) |
| *Spo11*<sup>-/-</sup> | 1692 | 335 | 895.8 (n=4) | 470 | 352 | 422.5 (n=4) | 387 | 95 | 265.2 (n=5) | 237 | 88 | 168.0 (n=4) |
| *Dmc1*<sup>-/-</sup> | 3554 | 487 | 1524.0 (n=4) | 620 | 89 | 350.0 (n=4) | 693 | 119 | 355.8 (n=5) | 365 | 51 | 137.0 (n=4) |
| *Hormad2*<sup>-/-</sup> | 4098 | 844 | 1836.3 (n=6) | 2311 | 253 | 1154.6 (n=5) | 1738 | 463 | 1048.2 (n=4) | 1473 | 205 | 597.8 (n=5) |
| *Spo11*<sup>-/-</sup> *Hormad2*<sup>-/-</sup> | 3045 | 2087 | 2535.8 (n=4) | 989 | 525 | 814.3 (n=4) | 1159 | 395 | 844.0 (n=4) | 1311 | 491 | 1022.0 (n=4) |
| *Dmc1*<sup>-/-</sup> *Hormad2*<sup>-/-</sup> | 764 | 302 | 563.3 (n=3) | 1191 | 369 | 819.7 (n=3) | 552 | 259 | 428.3 (n=4) | 401 | 156 | 289.0 (n=3) |

### Analysis of apoptotic oocyte frequency in meiotic mutants

By using this new method, we analyzed the oocyte apoptosis due to meiotic failure in *Spo11*^-/-^ and *Dmc1*^-/-^ ovaries during the perinatal period (Fig. 2). *Dmc1*^-/-^ oocytes having both DNA damage and extensive asynapsis showed a significant apoptosis increase from 17 dpc to 1dpp compared to wild-type oocytes (Fig. 2C and 2D). On the other hand, *Spo11*^-/-^ oocytes having extensive asynapsis without DNA damage showed a significant apoptosis increase only after birth (0 dpp and 1 dpp in Fig. 2D). These data indicate that DNA damage-dependent apoptosis might precede asynapsis-dependent apoptosis in mouse oocytes.

### HORMAD2-dependency of oocyte apoptosis due to meiotic deficiency

As HORMAD2 has been suggested to be involved in the elimination of meiotically aberrant oocytes (Rinaldi et al. 2017; Ravindranathan et al. 2022), we next examined HORMAD2-dependency of these meiotic apoptosis by generating double knockout mutants. Before that, we found that the lack of HORMAD2 significantly suppressed the apoptosis in wild-type oocytes only at 0 dpp (Fig. 3A), showing a presence of HORMAD2-dependent oocyte apoptosis in a physiological condition. HORMAD2 deficiency also suppressed a significant fraction of apoptosis in *Spo11*^-/-^ oocytes at 0 and 1 dpp (Fig. 3B) but not significantly in *Dmc1*^-/-^ oocytes at all ages (Fig. 3C). These results were consistent with the previous results showing a different HORMAD2-dependency of oocyte loss in SPO11- and DMC1-deficient oocytes (Kogo et al. 2012; Wojtasz et al. 2012; Rinaldi et al. 2017).

### Different timing of HORMAD2-dependent and -independent oocyte apoptosis

The time course of HORMAD2-independent and -dependent oocyte apoptosis was visualized separately by calculating the frequency of HORMAD2-dependent apoptosis in wild-type, *Spo11*^-/-^, and *Dmc1*^-/-^ oocytes (Fig. 4). The HORMAD2-independent apoptosis in DMC1-deficient oocytes (= the apoptosis in *Dmc1*^-/-^ *Hormad2*^-/-^ oocytes) was always significant from 17 dpc to 1dpp, peaking at 0 dpp, compared to that in wild-type oocytes (= the apoptosis in *Hormad2*^-/-^ oocytes) (Fig. 4A). Interestingly, HORMAD2-independent apoptosis in SPO11-deficient oocytes (= the apoptosis in *Spo11*^-/-^ *Hormad2*^-/-^ oocytes) was also significant at 0 dpp (p = 0.027), which might be caused by the SPO11-independent DSBs reported in late zygotene *Spo11*^-/-^ oocytes (Carofiglio et al. 2013). In contrast to that, the HORMAD2-dependent apoptosis was significant only in postnatal SPO11-deficient oocytes, especially at 1 dpp (Fig. 4B). In DMC1-deficient oocytes, HORMAD2-dependent apoptosis showed an increasing tendency at 1 dpp although not statistically significant (p = 0.052). These data clearly showed different kinetics between HORMAD2-independent and -dependent oocyte apoptosis, possibly consistent with the dual-checkpoint model for DNA damage and asynapsis (Ravindranathan et al. 2022).

**Figure 4.**
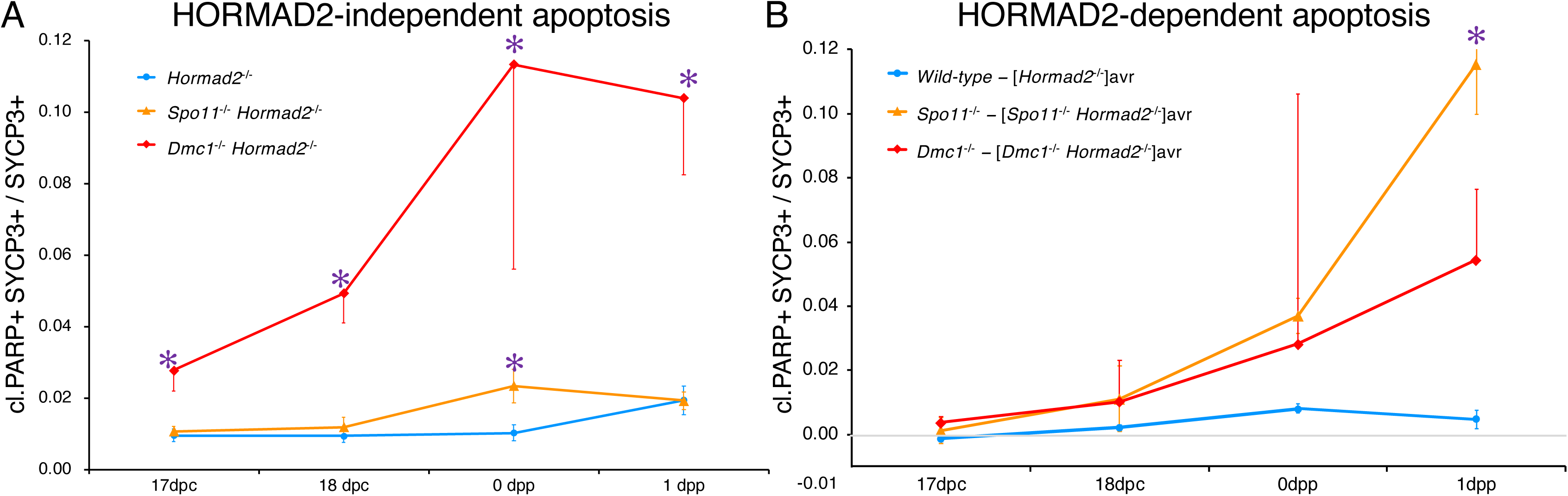
Time courses of HORMAD2-independent and -dependent apoptosis in ovaries having meiotic defects. Time courses of HORMAD2-independent (A) and -dependent (B) apoptosis frequencies in wild-type (blue), *Spo11*^-/-^ (orange), and *Dmc1*^-/-^ (red) perinatal mouse ovaries were shown in line graphs as mean ± SEM (n = 3-6). HORMAD2-dependent apoptosis was calculated by subtracting the average of corresponding HORMAD2-independent apoptosis frequencies from each apoptosis frequency in wild-type, *Spo11*^-/-^, and *Dmc1*^-/-^ ovaries. The data used were the same as those in Fig. 3. Statistical analysis was performed using the Kruskal-Wallis test, followed by the Steel’s test to compare the mutant groups (*Spo11*^-/-^ and *Dmc1*^-/-^) with the control group (wild-type). * p < 0.05. The HORMAD2-dependent apoptosis frequency in *Spo11*^-/-^ ovary was not statistically significant among the three groups due to the large variation in *Dmc1*^-/-^ ovaries at 0 dpp. However, it was significant when compared only with wild-type (p = 0.027).

### Future improvement points of the current method

In the present study, we employed SYCP3 and cleaved PARP1 as markers for meiotic and apoptotic nuclei, respectively. SYCP3 is a good marker but detected only in limited (from leptotene to diplotene) substages of oocytes. To include additional oocyte substages for analyses, other meiotic markers, such as meiosis-specific cohesins, should be employed (Lee et al. 2003; Burkhardt et al. 2016). As to apoptosis markers, it is unclear to what extent cleaved PARP1 could detect apoptotic oocytes with samples separated by 24 hours because cleaved PARP1 is likely expressed in a limited period of apoptotic process slightly preceding TUNEL staining (Huppertz et al. 1999; Pepling and Spradling 2001). Analyses of additional apoptosis markers will be helpful to understand the kinetics of meiotic apoptosis in more detail. When using other antibodies for immunostaining, buffers at basic pH, such as 20 mM Tris-HCl buffer (pH 9.0), would generally be adequate for heat-induced antigen retrieval (Yamashita 2007), although we used a citrate buffer (pH 6.0) that was optimal for our SYCP3 antibody.

As a specimen problem, the large center area of squashed ovaries was considerably thick and often unsuitable for immunofluorescent analysis. Although oocytes were fewer in this area than in the ovarian periphery, tissue-clearing techniques might be helpful in achieving more comprehensive analyses in squashed ovaries (Richardson and Lichtman 2015). In addition, erythrocytes sometimes showed intense autofluorescence in the squashed ovaries (Fig. 2 and Fig. S2, white arrowheads). Although the fluorescent signal of erythrocytes was distinguishable by their appearance, it might be required for other analyses to quench the erythrocyte autofluorescence by some treatments (Whittington and Wray 2017).

The present method has a disadvantage in correctly counting total oocytes, probably due to tissue squashing, as the analyzed oocyte numbers largely vary among specimens in the current protocol (Table 1). Whole-mount staining techniques would be a good substitute for that purpose, although some specialized fluorescent microscopes are required for observation.

In conclusion, the present method is easy, possibly applicable to a wide variety of antibodies, and can analyze enough oocytes to detect and analyze relatively infrequent events, facilitating detailed immunohistochemical analyses of fetal and perinatal mouse ovaries.

## Acknowledgements

We thank Dr. Yoko Shimizu and Dr. Yuka Kokubo-Kikuchi for their cell counting analysis in a non-involved position, and the staff at the Bioresource Center and the Laboratory for Analytical Instruments, Gunma University Graduate School of Medicine, for their technical support. In addition, we thank Ms. Mutsumi Shimoda for her assistance. We used research equipment shared in the MEXT Project for promoting public utilization of advanced research infrastructure (Program for supporting the introduction of the new sharing system) Grant Number JPMXS0420600120.

## Author contributions

Conceptualization: H.K.; Methodology: H.K., Y.T.; Validation: A.I-K., H.Y., M.I.; Formal analysis: H.K.; Investigation: H.K.; Resources: H.K., T.M.; Writing-original draft: H.K.; Writing-review & editing: H.K., Y.T., A.I-K., H.Y., M.I., T.M.; Visualization: H.K.; Supervision: T.M.; Project administration: H.K., T.M.; Funding acquisition: H.K.

## Data availability

No datasets were generated or analyzed during the current study.

---

**Figure S1.**
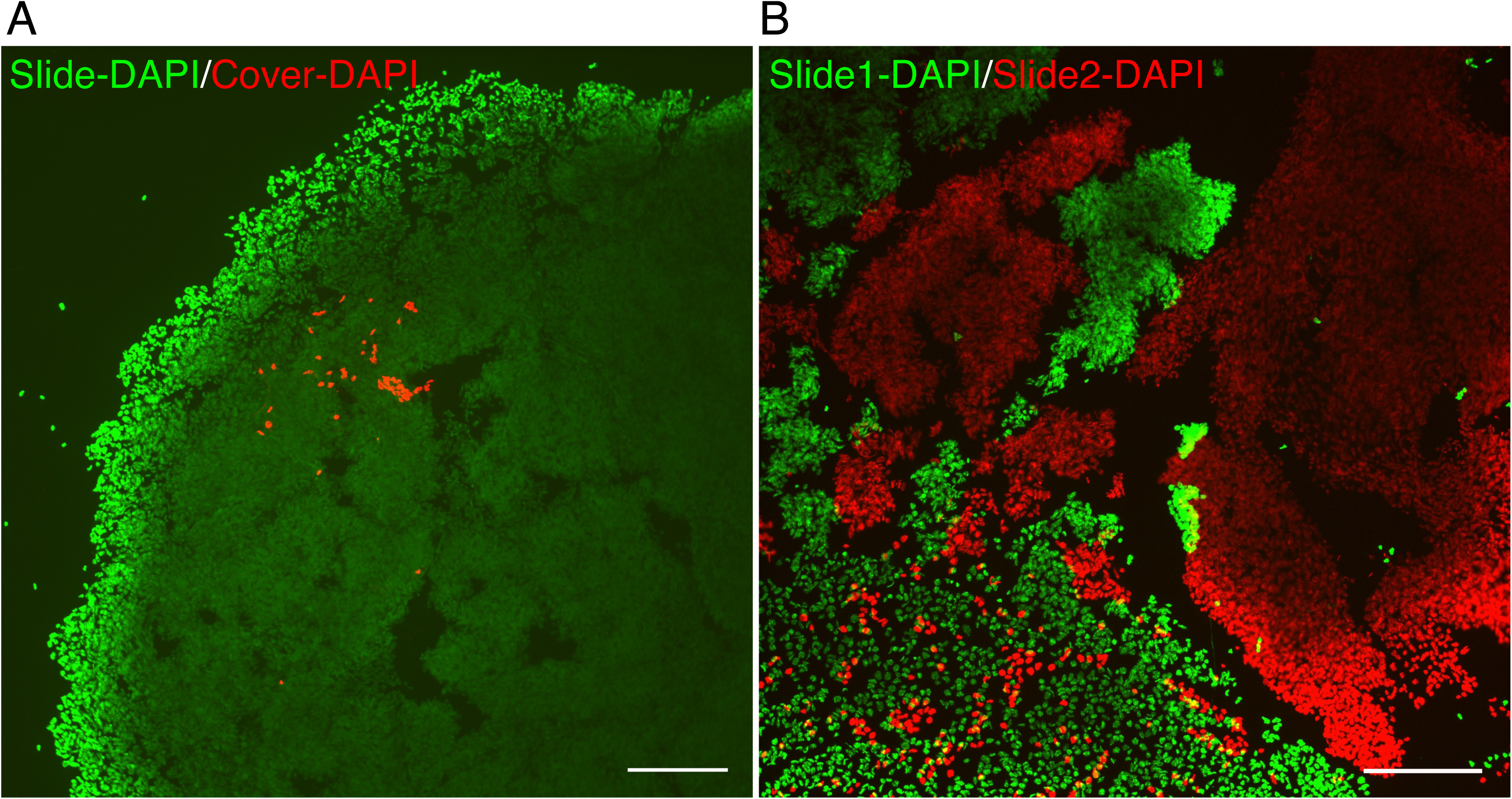
Evaluation of ovarian tissue adhesion to coverslips. Representative DAPI fluorescence images of squashed perinatal mouse ovaries using a silane-coated glass slide and a non-treated coverslip (A) or two silane-coated glass slides (B). Fluorescence images were taken separately, pseudo-colored in green (a glass slide in A and B) and in red (a coverslip in A and the other glass slide in B), and merged with the mirror image of the other to reproduce the original position of two surfaces facing each other. In contrast to the case using a non-treated coverslip (A), ovarian tissues adhered to both glass slides and separated randomly into pieces when ovaries were squashed by using two silane-coated glass slides (B). Scale bars are 200 µm.

**Figure S2.**
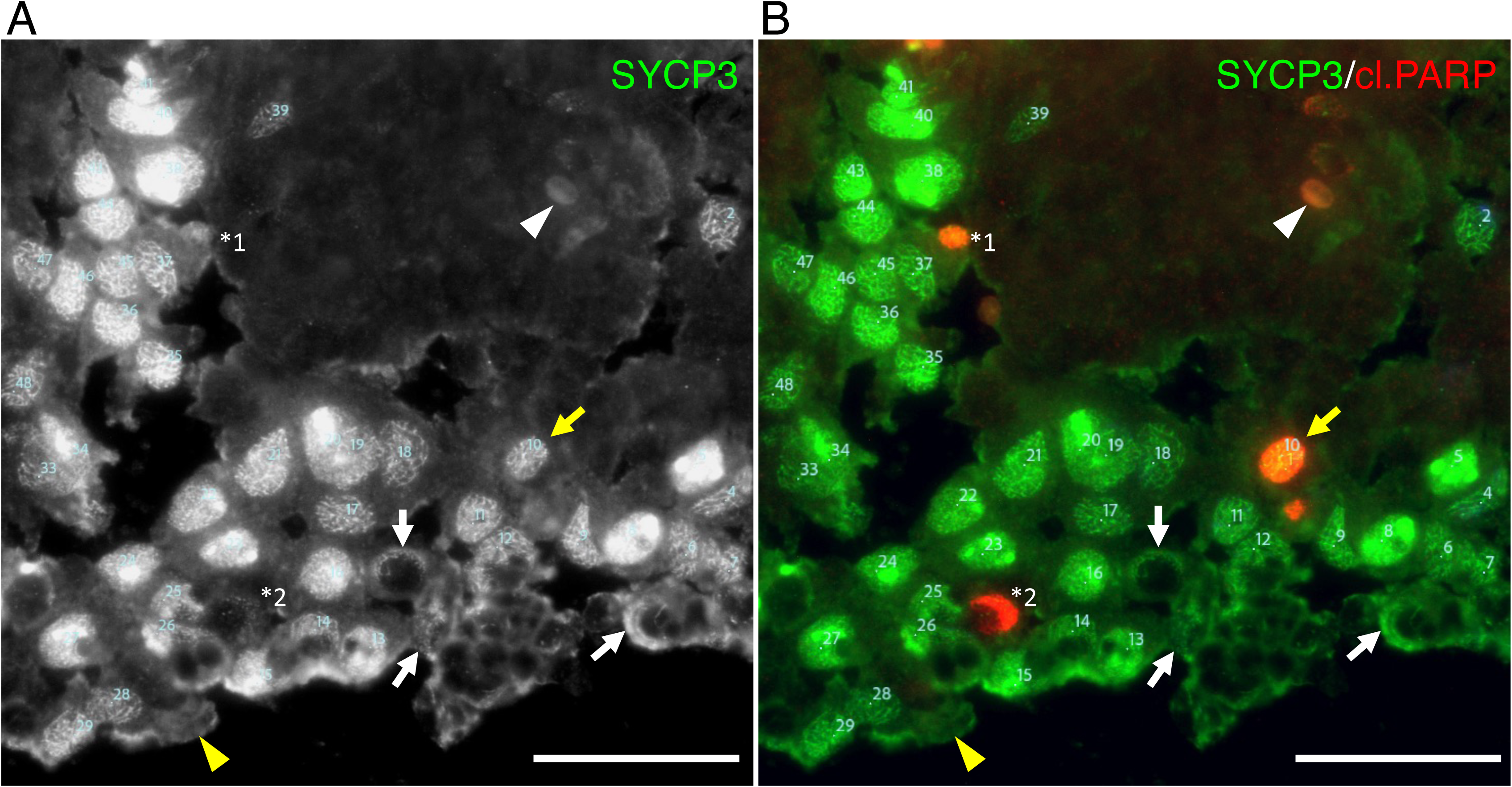
Analysis of meiotic oocyte apoptosis by manual counting. Example images of manual counting were shown (wild-type, 0 dpp). Firstly, the counter put sequential numbers on SYCP3-positive nuclei (numbered in light blue) in monochrome images using Photoshop’s counting tool (A). Diffuse signals of SYCP3, which were sometimes observed on erythrocytes (white arrowhead) and unidentified cells (yellow arrowhead), were omitted. Some SYCP3-positive nuclei were severely deformed (arrows) and also omitted from the analysis. After that, SYCP3- and cleaved PARP1-double positive nuclei were counted (numbered 1 in yellow on the nucleus numbered 10 in light blue, yellow arrow) in merged images where SYCP3 and cleaved PARP1 (cl.PARP) were pseudo-colored in green and red, respectively (B). In this case, cleaved PARP1-positive structures (*1 and *2) were possibly apoptotic oocyte nuclei but omitted due to the relatively small size (*1) and the weak signal of SYCP3 (*2), respectively. Scale bars are 50 µm.

## References

1. Baudat F, Manova K, Yuen JP, et al (2000) Chromosome Synapsis Defects and Sexually Dimorphic Meiotic Progression in Mice Lacking Spo11. Molecular Cell 6:989–998. 10.1016/S1097-2765(00)00098-8

2. Bolcun-Filas E, Schimenti JC (2012) Chapter Five - Genetics of Meiosis and Recombination in Mice. In: Jeon KW (ed) International Review of Cell and Molecular Biology. Academic Press, pp 179–227. 10.1016/B978-0-12-394309-5.00005-5

3. Bristol-Gould SK, Kreeger PK, Selkirk CG, et al (2006) Postnatal regulation of germ cells by activin: The establishment of the initial follicle pool. Developmental Biology 298:132–148. 10.1016/j.ydbio.2006.06.025

4. Burkhardt S, Borsos M, Szydlowska A, et al (2016) Chromosome Cohesion Established by Rec8-Cohesin in Fetal Oocytes Is Maintained without Detectable Turnover in Oocytes Arrested for Months in Mice. Curr Biol 26:678–685. 10.1016/j.cub.2015.12.073

5. Carofiglio F, Inagaki A, Vries S de, et al (2013) SPO11-Independent DNA Repair Foci and Their Role in Meiotic Silencing. PLOS Genetics 9:e1003538. 10.1371/journal.pgen.1003538

6. Di Giacomo M, Barchi M, Baudat F, et al (2005) Distinct DNA-damage-dependent and -independent responses drive the loss of oocytes in recombination-defective mouse mutants. Proceedings of the National Academy of Sciences 102:737–742. 10.1073/pnas.0406212102

7. Findlay JK, Hutt KJ, Hickey M, Anderson RA (2015) How Is the Number of Primordial Follicles in the Ovarian Reserve Established? Biol Reprod 93:111. 10.1095/biolreprod.115.133652

8. Huppertz B, Frank H-G, Kaufmann P (1999) The apoptosis cascade — morphological and immunohistochemical methods for its visualization. Anat Embryol 200:1–18. 10.1007/s004290050254

9. Kanda Y (2013) Investigation of the freely available easy-to-use software ‘EZR’ for medical statistics. Bone Marrow Transplant 48:452–458. 10.1038/bmt.2012.244

10. Kaur S, Kurokawa M (2023) Regulation of Oocyte Apoptosis: A View from Gene Knockout Mice. Int J Mol Sci 24:1345. 10.3390/ijms24021345

11. Kogo H, Tsutsumi M, Inagaki H, et al (2012) HORMAD2 is essential for synapsis surveillance during meiotic prophase via the recruitment of ATR activity. Genes to Cells 17:897–912. 10.1111/gtc.12005

12. Lee J, Iwai T, Yokota T, Yamashita M (2003) Temporally and spatially selective loss of Rec8 protein from meiotic chromosomes during mammalian meiosis. Journal of Cell Science 116:2781–2790. 10.1242/jcs.00495

13. McClellan KA, Gosden R, Taketo T (2003) Continuous loss of oocytes throughout meiotic prophase in the normal mouse ovary. Developmental Biology 258:334–348. 10.1016/S0012-1606(03)00132-5

14. Morita Y, Tilly JL (1999) Oocyte Apoptosis: Like Sand through an Hourglass. Developmental Biology 213:1–17. 10.1006/dbio.1999.9344

15. Morohaku K, Hirao Y, Obata Y (2017) Differentiation of Mouse Primordial Germ Cells into Functional Oocytes In Vitro. Ann Biomed Eng 45:1608–1619. 10.1007/s10439-017-1815-7

16. Niu W, Spradling AC (2022) Mouse oocytes develop in cysts with the help of nurse cells. Cell 185:2576–2590.e12. 10.1016/j.cell.2022.05.001

17. Page J, Suja JA, Santos JL, Rufas JS (1998) Squash procedure for protein immunolocalization in meiotic cells. Chromosome Res 6:639–642. 10.1023/a:1009209628300

18. Pepling ME (2006) From primordial germ cell to primordial follicle: mammalian female germ cell development. genesis 44:622–632. 10.1002/dvg.20258

19. Pepling ME, Spradling AC (2001) Mouse Ovarian Germ Cell Cysts Undergo Programmed Breakdown to Form Primordial Follicles. Developmental Biology 234:339–351. 10.1006/dbio.2001.0269

20. Pittman DL, Cobb J, Schimenti KJ, et al (1998) Meiotic Prophase Arrest with Failure of Chromosome Synapsis in Mice Deficient for Dmc1, a Germline-Specific RecA Homolog. Molecular Cell 1:697–705. 10.1016/S1097-2765(00)80069-6

21. Ravindranathan R, Raveendran K, Papanikos F, et al (2022) Chromosomal synapsis defects can trigger oocyte apoptosis without elevating numbers of persistent DNA breaks above wild-type levels. Nucleic Acids Research 50:5617–5634. 10.1093/nar/gkac355

22. Richardson DS, Lichtman JW (2015) Clarifying Tissue Clearing. Cell 162:246–257. 10.1016/j.cell.2015.06.067

23. Rinaldi VD, Bolcun-Filas E, Kogo H, et al (2017) The DNA Damage Checkpoint Eliminates Mouse Oocytes with Chromosome Synapsis Failure. Molecular Cell 67:1026–1036.e2. 10.1016/j.molcel.2017.07.027

24. Rodrigues P, Limback D, McGinnis LK, et al (2009) Multiple mechanisms of germ cell loss in the perinatal mouse ovary. 10.1530/REP-08-0203

25. Subramanian VV, Hochwagen A (2014) The Meiotic Checkpoint Network: Step-by-Step through Meiotic Prophase. Cold Spring Harb Perspect Biol 6:a016675. 10.1101/cshperspect.a016675

26. Whittington NC, Wray S (2017) Suppression of Red Blood Cell Autofluorescence for Immunocytochemistry on Fixed Embryonic Mouse Tissue. Current Protocols in Neuroscience 81:2.28.1–2.28.12. 10.1002/cpns.35

27. Wojtasz L, Cloutier JM, Baumann M, et al (2012) Meiotic DNA double-strand breaks and chromosome asynapsis in mice are monitored by distinct HORMAD2-independent and -dependent mechanisms. Genes Dev 26:958–973. 10.1101/gad.187559.112

28. Yamashita S (2007) Heat-induced antigen retrieval: Mechanisms and application to histochemistry. Progress in Histochemistry and Cytochemistry 41:141–200. 10.1016/j.proghi.2006.09.001

